# Recent SIV-Macaque Trials: A Reassessment of Immune Priming and Varying Infectability in Repeated Low-dose Challenge Studies

**DOI:** 10.1101/2021.07.28.454270

**Authors:** John L. Spouge

## Abstract

Nowadays, most preclinical HIV treatment trials use a protocol of administering repeated low-dose challenges (RLCs) of simian immunodeficiency virus (SIV) to macaques. Statistical analyses of treatment efficacy under the RLC protocol need to consider two confounding hypotheses, both pertinent biologically to HIV: (1) the non-infecting challenges may immunize animals against SIV; and (2) the animals may vary in intrinsic infectability (“frailty”). To explore the two hypotheses, a previous study (Regoes 2012) assembled a database from 7 articles with SIV-macaque treatment trials. With two explicable exceptions, Regoes concluded that the control data did not support either confounding hypothesis. Recent SIV-macaque trials present opportunities to evaluate the conclusions’ robustness. Accordingly, the present article assembles from 24 articles an updated database containing net survival curves from both control and treatment arms in SIV-macaque treatment trials. Broad patterns of statistical significance (at *p*<0.05, uncorrected for multiple testing) made it difficult to dismiss the confounding hypotheses completely in the controls. Although statistical analysis has focused on defense against variable frailty, only one set of controls showed significant variable frailty, whereas many sets showed significant immunization. As trials progressed, changes in the probability of infection per challenge were significant in 8/28 trials (1/3 trials using oral challenges; 2/4 trials using vaginal challenges; and 5/21 trials using rectal challenges). The results suggest the possibility that vaginal challenges may immunize animals faster than rectal challenges, and they also bear on previous conclusions that repeated exposure to HIV without treatment may have no effect on infectability or may even reduce it.

**Author Summary:** Many preclinical trials of HIV treatments rely on repeatedly administering low-dose SIV challenges to macaques until infection occurs. The repeated low-dose protocol reuses macaques and is more sensitive to subtle therapeutic efficacies than a protocol administering a single large dose to each macaque. The animal reuse raises some pertinent biological questions, notably: (1) do macaques have intrinsically variable infectabilities? and (2) do the repeated SIV challenges immunize macaques against infection? A 2012 study collected a database of eight macaque trials, concluding that variable infectability and immunization were at most sporadic and readily explicable. I expanded the 2012 database to twenty-eight trials, discovering that the conclusions were not robust. Although only 1/28 SIV-macaque trials showed variable infectability, 7/28 showed immunization, with few ready explanations. Statistical analysis of SIV-macaque trials has focused on the confounding effects of variable infectability to the neglect of immunization, so the expanded database provides a rich empirical resource. The trials have general medical importance because they provide a model for analyzing animal trials of infectious disease therapies and other sparse trials, e.g., for breast cancer. My findings also indirectly suggest that repeated human exposure to HIV inconsistently immunizes and can foster either immune priming or tolerance.

## Introduction

Preclinical experiments play a fundamental role in determining treatments for an infectious disease. In a modern re-interpretation, some of Koch’s postulates specify reproducing the disease in an animal model and then reisolating the agent from an animal with signs resembling the human disease [1]. Historically, experimenters therefore tested vaccines or preventive treatment strategies against human-immunodeficiency virus (HIV) preclinically in non-human primates. They found that HIV infects in the chimpanzee animal model, although it does not consistently produce an AIDS-like disease.

Typically, preclinical HIV trials have control and treatment arms, and they infer therapeutic protection if treated animals are less easily infected than controls. In the 1980s, preclinical HIV trials were extremely sparing because of the cost of chimpanzees, e.g., one trial used 6 chimpanzees in a control arm and 2 chimpanzees in a treatment arm [2]. Such trials ensured infection of controls with a large viral challenge [3, 4]. The trials then applied a statistical test, e.g., the Fisher’s exact test [5], to the resulting 2×2 tables of counts of animals in control/treated vs infected/uninfected groups. Unfortunately, with so few counts, the single high-dose protocol provided scant statistical power for detecting subtle protection against infection. Accordingly, experimenters relied heavily on surrogates of protection to detect a treatment effect, e.g., an increase in virus-specific immune responses from a vaccine treatment, or a decrease in the steady-state viral load (a.k.a. the viral set-point). The surrogates are chosen to correlate with protection, although they are not necessarily causal.

Since 2007, chimpanzees are an endangered species no longer bred for research [6], requiring other animal models for HIV infection and AIDS. Mice can receive transplanted components of a human immune system [7], and when applicable, the humanized mouse model is cost-effective. The mouse model has, however, many serious limitations in cell number and volume, life span, integrated operation of an immune system, etc. [8]. By contrast, over 40 different simian immunodeficiency viruses (SIVs) infect primate species without such limitations [9], and some do produce AIDS-like illnesses. Presently, the rhesus macaque and its SIV provide the most common preclinical primate model of AIDS [10–33].

Although rhesus macaques provide less expensive preclinical data than chimpanzees, their cost is still severe. Excluding the historical controls used in recent trials [26, 29, 32], macaque trials have limited themselves on average to about 13 contemporary controls. Under such limitations, if the trials used the single high-dose protocol, they would remain seriously underpowered for detecting a subtle treatment effect [34].

Accordingly, preclinical HIV treatment trials have adopted a protocol using repeated low-dose challenges (RLCs) [10–33]. As Table 1 of [18] helpfully illustrates, typically each animal is challenged repeatedly at regular intervals with the same low viral dose, until either it is infected or the experiment finishes. Experimenters usually challenge weekly, because the anesthetized animal can then receive the current challenge after tests for infection from the previous week are initiated. To limit the number of challenges required to infect controls, the challenge dose often approximates the animal ID_50_ [4]. Some experiments challenge beyond infection (e.g., [10]), but post-infection challenges are irrelevant to present purposes, and we ignore them.

**Table 1:**
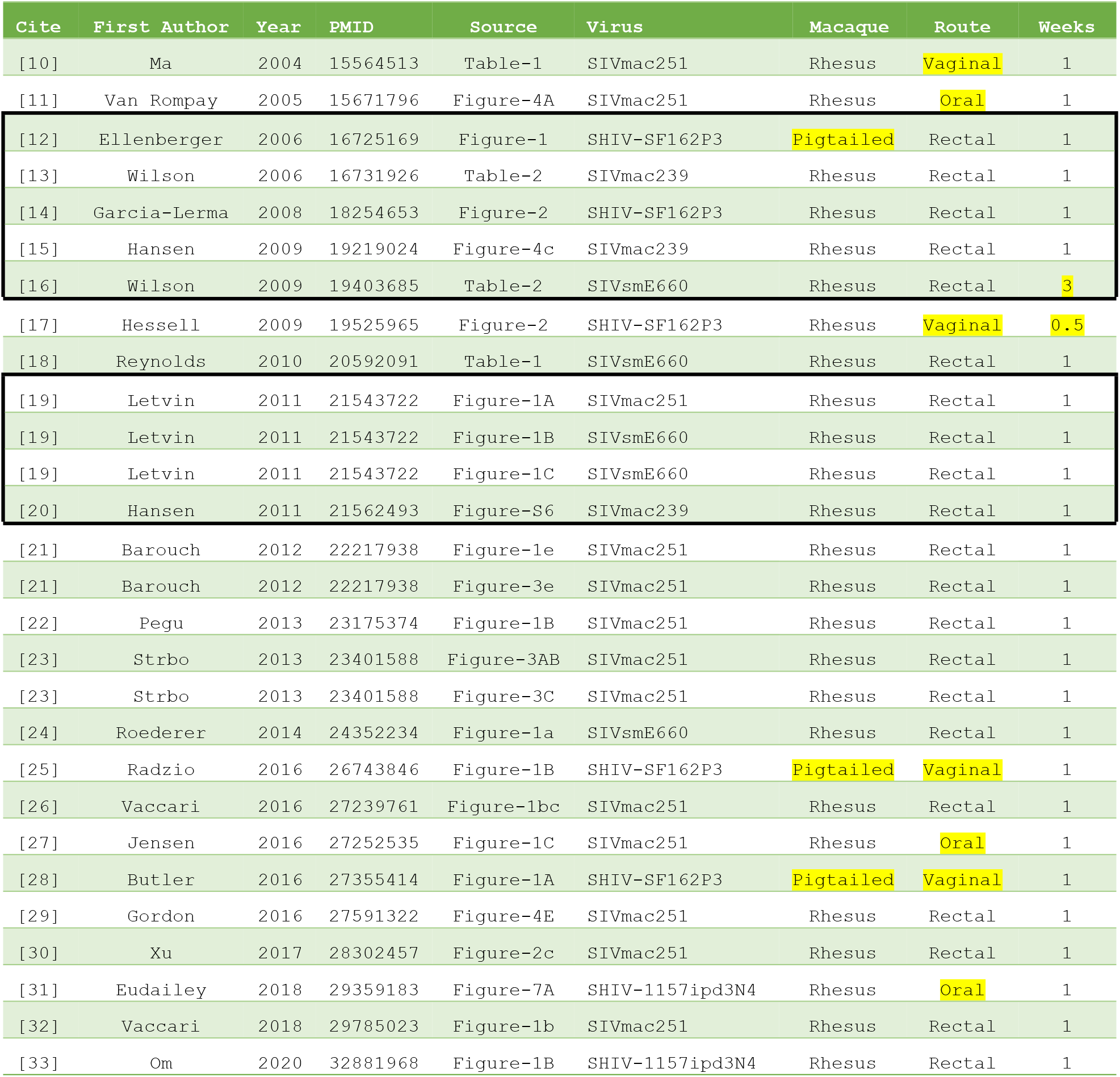
Metadata for the SIV-Macaque database.

| Cite | First Author | Year | PMID | Source | Virus | Macaque | Route | Weeks |
| --- | --- | --- | --- | --- | --- | --- | --- | --- |
| [10] | Ma | 2004 | 15564513 | Table-1 | SIVmac251 | Rhesus | Vaginal | 1 |
| [11] | Van Rompay | 2005 | 15671796 | Figure-4A | SIVmac251 | Rhesus | Oral | 1 |
| [12] | Ellenberger | 2006 | 16725169 | Figure-1 | SHIV-SF162P3 | Pigtailed | Rectal | 1 |
| [13] | Wilson | 2006 | 16731926 | Table-2 | SIVmac239 | Rhesus | Rectal | 1 |
| [14] | Garcia-Lerma | 2008 | 18254653 | Figure-2 | SHIV-SF162P3 | Rhesus | Rectal | 1 |
| [15] | Hansen | 2009 | 19219024 | Figure-4c | SIVmac239 | Rhesus | Rectal | 1 |
| [16] | Wilson | 2009 | 19403685 | Table-2 | SIVsmE660 | Rhesus | Rectal | 3 |
| [17] | Hessell | 2009 | 19525965 | Figure-2 | SHIV-SF162P3 | Rhesus | Vaginal | 0.5 |
| [18] | Reynolds | 2010 | 20592091 | Table-1 | SIVsmE660 | Rhesus | Rectal | 1 |
| [19] | Letvin | 2011 | 21543722 | Figure-1A | SIVmac251 | Rhesus | Rectal | 1 |
| [19] | Letvin | 2011 | 21543722 | Figure-1B | SIVsmE660 | Rhesus | Rectal | 1 |
| [19] | Letvin | 2011 | 21543722 | Figure-1C | SIVsmE660 | Rhesus | Rectal | 1 |
| [20] | Hansen | 2011 | 21562493 | Figure-S6 | SIVmac239 | Rhesus | Rectal | 1 |
| [21] | Barouch | 2012 | 22217938 | Figure-1e | SIVmac251 | Rhesus | Rectal | 1 |
| [21] | Barouch | 2012 | 22217938 | Figure-3e | SIVmac251 | Rhesus | Rectal | 1 |
| [22] | Pegu | 2013 | 23175374 | Figure-1B | SIVmac251 | Rhesus | Rectal | 1 |
| [23] | Strbo | 2013 | 23401588 | Figure-3AB | SIVmac251 | Rhesus | Rectal | 1 |
| [23] | Strbo | 2013 | 23401588 | Figure-3C | SIVmac251 | Rhesus | Rectal | 1 |
| [24] | Roederer | 2014 | 24352234 | Figure-1a | SIVsmE660 | Rhesus | Rectal | 1 |
| [25] | Radzio | 2016 | 26743846 | Figure-1B | SHIV-SF162P3 | Pigtailed | Vaginal | 1 |
| [26] | Vaccari | 2016 | 27239761 | Figure-1bc | SIVmac251 | Rhesus | Rectal | 1 |
| [27] | Jensen | 2016 | 27252535 | Figure-1C | SIVmac251 | Rhesus | Oral | 1 |
| [28] | Butler | 2016 | 27355414 | Figure-1A | SHIV-SF162P3 | Pigtailed | Vaginal | 1 |
| [29] | Gordon | 2016 | 27591322 | Figure-4E | SIVmac251 | Rhesus | Rectal | 1 |
| [30] | Xu | 2017 | 28302457 | Figure-2c | SIVmac251 | Rhesus | Rectal | 1 |
| [31] | Eudailey | 2018 | 29359183 | Figure-7A | SHIV-1157ipd3N4 | Rhesus | Oral | 1 |
| [32] | Vaccari | 2018 | 29785023 | Figure-1b | SIVmac251 | Rhesus | Rectal | 1 |
| [33] | Om | 2020 | 32881968 | Figure-1B | SHIV-1157ipd3N4 | Rhesus | Rectal | 1 |

The RLC protocol has several advantages [35], both biological and statistical [36]. First, RLCs mimic human exposure to HIV, making the animal studies conceptually closer to subsequent human studies. Second, they can in principle detect immunization of controls statistically during repeated challenge, as follows. RLCs yield net survival curves for each control or treatment arm (e.g., Figure 3 of [18]), where the curves count the uninfected animals before and after each challenge. Non-infecting challenges might gradually immunize animals, but statistical analysis of the fraction of controls infected after each challenge can detect, e.g., systematic decreases in the fraction over time. In effect, the statistical analysis defines “infectability” as a probability of infection, with “immunization” as the corresponding change in infectability. To paraphrase [35], the statistical definition of immunization presumes neither knowledge nor measurement of the immune surrogates of protection, but instead specifies what matters epidemiologically. In contrast, biological studies examining laboratory surrogates of protection (e.g., [2]) implicitly rely on a strong but unproved assumption, that the laboratory surrogates are causally linked to immune protection [37]. Though limited in other ways, a statistical definition of immunization deftly avoids implicit assumptions about causality.

Third, RLCs can also expose intrinsic differences in infectability between subpopulations of animals, their “frailties” in the terminology of survival analysis [35]. Immune surrogates have demonstrated some specific subpopulation differences, e.g., the peak plasma RNA in SIVsmE660 infection is about 10-fold reduced in rhesus macaques with the *Mamu*-*A*01* class I allele [19]. On one hand, therefore, a study examining immune surrogates can draw specific biological conclusions. On the other hand, a statistical analysis of infectability has the complementary strength of generality, because it may detect immunization or variable frailty regardless of specific biological mechanisms like alleles or restriction factors [38]. Unfortunately, because of a phenomenon known as “heterogeneity’s ruse” [39] (see also the Discussion and the SI), statistical analysis of a net survival curve does not dependably distinguish variable frailty from immunization. Further biological analysis is often still required to disentangle their effects [35].

The RLC protocol has already demonstrated substantial advantages over the single high-dose protocol [36], but some advantages may remain unrealized. Presently, statistical analyses of many SIV-macaque trials use a standard survival analysis with the logrank test [22–24, 26, 29, 32]. The standard logrank test requires a large-sample Gaussian approximation, whereas the sparsity of SIV-macaque data suggests the need for an exact test. The development of an exact test is eased if immunization, variable frailty, or other confounding effects do not occur in SIV-macaque trials, because then every challenge in an RLC protocol presents the same hazard of infection to every control animal [40, 41]. In 2012, therefore, Regoes collected a database of 7 articles with SIV-macaque treatment trials, concluding that most did not display immunization or variable frailty [35].

In 2015, Nolen et al [42] pointed out that a Fisher exact test is valid in a single high-dose challenge study, but not in a RLC study if it is applied to the total challenge counts in different arms of the trial. They therefore suggested exact permutation tests based on exchangeability of control and treatment animals under the null hypothesis of treatment having no effect. After a careful theoretical analysis, they applied the permutation tests, demonstrating that large-sample p-value approximations could yield misleading scientific conclusions in small trials. In passing, however, observe that the single Ellenberger (2006) trial [17] mentioned explicitly in Section 5 of Nolen et al [42] was mildly atypical (see comments in the Results), and that the only confounding hypotheses Nolen et al [42] simulated involved variable frailty. These two observations partially motivated the present article by suggesting that statistical analysis might profit from a comprehensive database of SIV-macaque trials.

Although the present article notes some technical points, it does not improve substantially on Regoes’ statistical analysis of immunization and variable frailty [35]. New SIV-macaque trials contain many pertinent data, however. The present article therefore assembles a comprehensive updated database from 24 articles containing the net survival curves for control and treatment arms in SIV-macaque treatment trials. It has the purview of applying Regoes’ statistical analysis to a larger database to examine the robustness of his scientific conclusions.

The RLC protocol has regularities not available in clinical trials, notably compulsory follow up and a possibility that between observations, all hazards to controls may be roughly the same, i.e., all probabilities of infection per challenge are roughly equal. The database may therefore provide useful biological examples for applications with data more irregular than SIV-macaque treatment trials but still requiring the development of small-sample statistical tests, e.g., small studies of breast cancer therapies [43]. As Regoes also pointed out [35], the database also provides an animal model for exploring repeated low-dose exposure to HIV in humans without treatment.

The present article is structured as follows. The Materials and Methods section gives the criteria for trial data to enter our SIV-Macaque database of control results. The human effort in assembly of the database and its subsequent manipulation was demanding, so the Materials and Methods section discusses at length the various representations of data in the original articles, during digitization of survival curves, and for the survival analysis [44]. It then gives Regoes’ models and two additional full models (Geometric priming and Delta frailty), promoting continuity with Regoes’ statistical analysis [35] by using his term “priming models” for immunization. The Results section presents three tables pertinent to our SIV-Macaque database: (1) the metadata; (2) the count of controls uninfected after each trial’s maximum number of challenges, with estimates of the infection probability per challenge under the assumption that the probability is constant; and (3) p-values pertinent to the constancy of infection probabilities per challenge. The three tables are drawn from an aggregate table in the SI. Finally, the Discussion section explores the implications of the results.

## Results

The comprehensive Excel file in the SI yielded all rounded numbers in Tables 1–3.

Table 1 presents the metadata of the SIV-Macaque database. Each row corresponds to a single control arm and its trial. The columns indicate: (1) the citation in the present article; (2) its first author; (3) its year of publication; (4) its ID in PubMed; (5) the source within it for each set of controls (a citation may contain more than one trial and therefore more than one source of data); (6) the virus used to challenge; (7) the species of macaque challenged; (8) the challenge route; and (9) the number of weeks between the regular challenges. In the final 3 columns, uncommon values are highlighted in yellow, e.g., the Ellenberger (2006) trial [12] used SHIV-SF162P3 to challenge pigtailed macaques rectally every week, the Wilson (2009) trial [16] used SIVsmE660 to challenge rhesus macaques rectally every 3 weeks, etc.

The first 5 columns of Table 1 appear in all tables, so the reader can recall references and still find them immediately and unambiguously in the literature. In all tables, the rows are ordered on the PubMed ID in Column 4; and the heavy borders enclose the Regoes’ dataset [35].

Table 2 focuses on results pertinent to the Constant Hazard reduced model, which assumes that every challenge has a constant probability of infection. Columns after Column 5 indicate: (6) the number *n* of control animals; (7) the number of uninfected controls remaining after RLCs, with non-0 values highlighted in yellow; (8) the number *m* of RLCs that uninfected controls survived; (9) the MLE 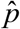 of the probability *p* of infection in the reduced model; and the (10) lower and (11) upper endpoint of the corresponding 95% confidence interval for *p*. Thus, Table 2 indicates for macaque trials the potential sparsity of the control arm, controls remaining uninfected after RLCs, and an approximation to the infection probability *p* of each challenge.

**Table 2:** Constant Hazard reduced model for controls in the SIV-Macaque database.

| Cite | First Author | Year | PMID | Source | n | Not<br>inf | m | p_hat | p_lo | p_hi |
| --- | --- | --- | --- | --- | --- | --- | --- | --- | --- | --- |
| [10] | Ma | 2004 | 15564513 | Table-1 | 8 | 0 | 5 | 0.364 | 0.186 | 0.572 |
| [11] | Van Rompay | 2005 | 15671796 | Figure-4A | 5 | 0 | 25 | 0.312 | 0.125 | 0.555 |
| [12] | Ellenberger | 2006 | 16725169 | Figure-1 | 14 | 1 | 13 | 0.206 | 0.119 | 0.317 |
| [13] | Wilson | 2006 | 16731926 | Table-2 | 8 | 0 | 8 | 0.216 | 0.105 | 0.365 |
| [14] | Garcia-Lerma | 2008 | 18254653 | Figure-2 | 18 | 1 | 14 | 0.205 | 0.128 | 0.300 |
| [15] | Hansen | 2009 | 19219024 | Figure-4c | 15 | 0 | 12 | 0.242 | 0.147 | 0.358 |
| [16] | Wilson | 2009 | 19403685 | Table-2 | 8 | 2 | 5 | 0.250 | 0.108 | 0.443 |
| [17] | Hessell | 2009 | 19525965 | Figure-2 | 4 | 0 | 40 | 0.400 | 0.146 | 0.700 |
| [18] | Reynolds | 2010 | 20592091 | Table-1 | 8 | 1 | 10 | 0.318 | 0.151 | 0.525 |
| [19] | Letvin | 2011 | 21543722 | Figure-1A | 20 | 0 | 5 | 0.500 | 0.349 | 0.651 |
| [19] | Letvin | 2011 | 21543722 | Figure-1B | 25 | 3 | 12 | 0.200 | 0.133 | 0.281 |
| [19] | Letvin | 2011 | 21543722 | Figure-1C | 20 | 5 | 12 | 0.124 | 0.073 | 0.190 |
| [20] | Hansen | 2011 | 21562493 | Figure-S6 | 28 | 1 | 25 | 0.156 | 0.107 | 0.215 |
| [21] | Barouch | 2012 | 22217938 | Figure-1e | 8 | 0 | 6 | 0.727 | 0.435 | 0.924 |
| [21] | Barouch | 2012 | 22217938 | Figure-3e | 8 | 0 | 6 | 0.533 | 0.291 | 0.765 |
| [22] | Pegu | 2013 | 23175374 | Figure-1B | 11 | 0 | 5 | 0.333 | 0.189 | 0.502 |
| [23] | Strbo | 2013 | 23401588 | Figure-3AB | 12 | 0 | 10 | 0.480 | 0.293 | 0.671 |
| [23] | Strbo | 2013 | 23401588 | Figure-3C | 12 | 0 | 6 | 0.324 | 0.189 | 0.483 |
| [24] | Roederer | 2014 | 24352234 | Figure-1a | 20 | 0 | 12 | 0.339 | 0.227 | 0.465 |
| [25] | Radzio | 2016 | 26743846 | Figure-1B | 5 | 0 | 16 | 0.313 | 0.125 | 0.555 |
| [26] | Vaccari | 2016 | 27239761 | Figure-1bc | 47 | 0 | 10 | 0.367 | 0.287 | 0.453 |
| [27] | Jensen | 2016 | 27252535 | Figure-1C | 6 | 0 | 9 | 0.240 | 0.103 | 0.428 |
| [28] | Butler | 2016 | 27355414 | Figure-1A | 11 | 5 | 20 | 0.045 | 0.018 | 0.089 |
| [29] | Gordon | 2016 | 27591322 | Figure-4E | 47 | 0 | 10 | 0.367 | 0.287 | 0.453 |
| [30] | Xu | 2017 | 28302457 | Figure-2c | 6 | 2 | 5 | 0.235 | 0.080 | 0.465 |
| [31] | Eudailey | 2018 | 29359183 | Figure-7A | 7 | 1 | 8 | 0.176 | 0.074 | 0.326 |
| [32] | Vaccari | 2018 | 29785023 | Figure-1b | 53 | 0 | 10 | 0.414 | 0.331 | 0.500 |
| [33] | Om | 2020 | 32881968 | Figure-1B | 16 | 7 | 3 | 0.237 | 0.122 | 0.386 |

For the Hansen (2009) trial [15], Regoes [35] gives the control number as *n* =15, whereas the legend of Figure-4c, the source, gives *n* =16, yielding minor numerical discrepancies of no scientific substance. In the Hansen (2011) trial [20], the single uninfected control eventually showed signs of infection, but long after the 1-week window for establishing infection during the trial. Its designation as “uninfected” is therefore somewhere arbitrary but at least consistent with the intent of the RLC protocol. Some controls were historical compared to treated animals [26, 29, 32]. Both the Vaccari (2016) trial [26] and the Gordon (2016) trial [29] used 23 concurrent and 24 historical controls. The two control datasets yielded the same net survival curve. To make the presentation consistent, the statistical analysis appears twice in all tables, once for each trial. The Vaccari (2018) trial [32] used 6 concurrent and 47=23+24 historical controls.

Table 3 focuses on various full models nesting the Constant Hazard reduced model, and it displays the p-values (uncorrected for multiple testing) from the corresponding LRTs. Columns after the 5-th indicate the p-values for the following full models: (6) Arithmetic Priming (AP); (7) Geometric Priming (GP); (8) Step Priming with *l*_*step*_ =1 (SP1); (9) Step Priming with *l*_*step*_ =2 (SP2); (10) Step Priming with *l*_*step*_ =3 (SP3); (11) Beta Frailty (BF); and (12) Delta Frailty (DF). In the final 7 columns, any p-value less than 0.05 is highlighted in yellow.

**Table 3:** Likelihood ratio test (LRT) p-values for the SIV-Macaque database.

| Cite | First Author | Year | PMID | Source | AP | GP | SP1 | SP2 | SP3 | BF | DF |
| --- | --- | --- | --- | --- | --- | --- | --- | --- | --- | --- | --- |
| [10] | Ma | 2004 | 15564513 | Table-1 | 0.084 | 0.060 | 0.395 | 0.054 | 0.250 | 1.000 | 1.000 |
| [11] | Van Rompay | 2005 | 15671796 | Figure-4A | 0.390 | 0.229 | 0.099 | 0.045 | 0.097 | 0.110 | 1.000 |
| [12] | Ellenberger | 2006 | 16725169 | Figure-1 | 0.446 | 0.406 | 0.492 | 0.098 | 0.131 | 0.380 | 0.600 |
| [13] | Wilson | 2006 | 16731926 | Table-2 | 0.017 | 0.024 | 0.035 | 0.036 | 0.428 | 1.000 | 1.000 |
| [14] | Garcia-Lerma | 2008 | 18254653 | Figure-2 | 0.425 | 0.371 | 0.837 | 0.176 | 0.155 | 0.321 | 0.665 |
| [15] | Hansen | 2009 | 19219024 | Figure-4c | 0.856 | 0.834 | 0.799 | 0.307 | 0.992 | 1.000 | 1.000 |
| [16] | Wilson | 2009 | 19403685 | Table-2 | 1.000 | 1.000 | 0.297 | 0.807 | 0.768 | 1.000 | 0.825 |
| [17] | Hessell | 2009 | 19525965 | Figure-2 | 0.048 | 0.103 | 0.016 | 0.749 | 0.158 | 1.000 | 1.000 |
| [18] | Reynolds | 2010 | 20592091 | Table-1 | 0.020 | 0.028 | 0.170 | 0.036 | 0.124 | 0.043 | 0.023 |
| [19] | Letvin | 2011 | 21543722 | Figure-1A | 0.810 | 0.769 | 0.204 | 0.489 | 0.677 | 0.696 | 1.000 |
| [19] | Letvin | 2011 | 21543722 | Figure-1B | 0.101 | 0.076 | 0.234 | 0.027 | 0.033 | 0.086 | 0.119 |
| [19] | Letvin | 2011 | 21543722 | Figure-1C | 0.214 | 0.222 | 0.705 | 0.407 | 0.706 | 0.238 | 0.217 |
| [20] | Hansen | 2011 | 21562493 | Figure-S6 | 0.269 | 0.264 | 0.371 | 0.319 | 0.868 | 0.275 | 0.320 |
| [21] | Barouch | 2012 | 22217938 | Figure-1e | 0.910 | 0.898 | 0.785 | 0.412 | 1.000 | 1.000 | 1.000 |
| [21] | Barouch | 2012 | 22217938 | Figure-3e | 0.504 | 0.454 | 0.782 | 0.601 | 0.251 | 1.000 | 1.000 |
| [22] | Pegu | 2013 | 23175374 | Figure-1B | 0.006 | 0.007 | 0.025 | 0.022 | 0.183 | 1.000 | 1.000 |
| [23] | Strbo | 2013 | 23401588 | Figure-3AB | 0.877 | 0.869 | 0.847 | 0.568 | 0.584 | 1.000 | 1.000 |
| [23] | Strbo | 2013 | 23401588 | Figure-3C | 0.003 | 0.003 | 0.020 | 0.012 | 0.058 | 1.000 | 1.000 |
| [24] | Roederer | 2014 | 24352234 | Figure-1a | 0.966 | 0.962 | 0.898 | 0.652 | 0.506 | 1.000 | 1.000 |
| [25] | Radzio | 2016 | 26743846 | Figure-1B | 0.096 | 0.089 | 0.030 | 0.889 | 0.361 | 1.000 | 1.000 |
| [26] | Vaccari | 2016 | 27239761 | Figure-1bc | 0.267 | 0.268 | 0.388 | 0.258 | 0.449 | 1.000 | 1.000 |
| [27] | Jensen | 2016 | 27252535 | Figure-1C | 0.655 | 0.557 | 0.549 | 0.569 | 0.910 | 1.000 | 1.000 |
| [28] | Butler | 2016 | 27355414 | Figure-1A | 0.052 | 0.133 | 0.315 | 0.993 | 0.600 | 0.224 | 0.099 |
| [29] | Gordon | 2016 | 27591322 | Figure-4E | 0.267 | 0.268 | 0.388 | 0.258 | 0.449 | 1.000 | 1.000 |
| [30] | Xu | 2017 | 28302457 | Figure-2c | 0.181 | 0.257 | 0.488 | 0.442 | 0.121 | 0.305 | 0.222 |
| [31] | Eudailey | 2018 | 29359183 | Figure-7A | 0.476 | 0.386 | 0.417 | 0.912 | 0.364 | 1.000 | 1.000 |
| [32] | Vaccari | 2018 | 29785023 | Figure-1b | 0.178 | 0.180 | 0.282 | 0.246 | 0.267 | 1.000 | 1.000 |
| [33] | Om | 2020 | 32881968 | Figure-1B | 0.738 | 0.757 | 0.538 | 0.906 | 1.000 | 1.000 | 1.000 |

Table 3 had 8/28 rows corresponding to trials displaying deviation from a constant infection probability per challenge at statistical significance *p*<0.05, uncorrected for multiple testing. The 8 trials showing deviation included 1/3 trials using oral challenges, 2/4 trials using vaginal challenges, and 5/21 using rectal challenges. For each full model of immune priming with a significant LRT, the second of its two MLE parameters (given in the SI) indicates whether the infection probability per challenge increased or decreased (tolerance or priming) as the trial progressed. The direction of change, increase or decrease, was different in different trials.

## Discussion

All p-values below are uncorrected for multiple testing.

In the tables, the rows inside the thick borders correspond to Regoes’ database [35]. The rows in Table 2 differ from Regoes’ Table 2 in: (1) the subdivision of data from the Letvin (2011) trial [19]; and (2) the number of controls in the Hansen (2009) trial [15]. Table 3 and Regoes’ Tables 3 and 4 have analogous differences. Otherwise, the present analysis agrees broadly within numerical error on its overlap with Regoes’ [35]. Regoes’ Tables 3 and 4 displayed only a few anomalous rows with *p*<0.05, consistent with his conclusion that a constant hazard of infection per challenge could explain most of his data [35]. The anomaly in the Wilson (2006) trial was ascribed to possible misidentification of infecting challenges; the anomaly in the Letvin (2011) Figure-1B trial, to the known biological effect of TRIM5 alleles on SIV infection (e.g., [52]).

The trials in Table 3 outside Regoes’ Tables 3 and 4 continue to display a broad pattern of anomalies with *p*<0.05, however, making the anomalies harder to dismiss. Perhaps some anomalies still reflect misidentification of the infecting challenges, but the misidentification then becomes a persistent feature of the SIV-macaque trials. Other anomalies may depend on trials using oral or vaginal challenge, which Regoes [35] excluded. The remaining anomalies may reflect unknown and therefore interesting biological effects. The persistent pattern of anomalies in Table 3 made Regoes’ extensive analysis of the power of the LRTs against alternatives unnecessary. By themselves, the anomalies clearly discourage any simplifying assumption of a constant hazard of infection per challenge in SIV-macaque trials, particularly for oral or vaginal challenges, particularly without specific evidence that supports the assumption’s use as an approximation.

The SIV-Macaque database and Table 3 noticeably enrich the empirical basis for the statistical analysis of sparse animal trials. Presently, permutation tests provide the leading candidates for exact tests. The power analysis of the permutation tests in [42], e.g., focused on confounding hypotheses of variable frailty, namely, the two frailty full models in the present article. The frailty models accurately reflect experimenters’ concerns about the confounding effects of, e.g., sex [33] or the *Mamu*-*A*01* class I allele [19]. Table 3 suggests, however, that a statistical exploration of the two confounding hypotheses should probably be more concerned about immunization than variable frailty in [42].

The pattern of anomalies in Table 3 also has a notable feature. Except for the Hessell (2009) trial [17] and the Radzio (2016) trial [25], both of which challenged vaginally, whenever a full model in Table 3 displayed *p*<0.05, the full model SP2 for step priming after two challenges in the same row also displayed *p*<0.05. Under a weekly challenge, the full model SP2 could be interpreted as changes in control immunity occurring after 2 weeks, e.g., the recommended period for full immunity after COVID-19 vaccination. The immune changes were inconsistent: neither immune priming nor immune tolerance occurred consistently (see end of the Results and the SI). For the full model SP2, e.g., 3/6 changes indicated increased probability of infection per challenge (tolerance), rather than priming. Typically, however, net survival curves alone cannot distinguish variable frailty from immunization (“heterogeneity’s ruse” [39]). The inconsistent immune interpretation of the SP2 full model does, however, bear on previous biological conclusions that repeated exposure without treatment may have no effect [35] or may even reduce [53] human infectability in HIV acquisition.

The exceptions above, the Hessell (2009) trial [17] and the Radzio (2016) trial [25], used vaginal challenge. Neither had full model SP2 displaying *p*<0.05, but instead had full model SP1 for step priming after one challenge displaying *p*<0.05, suggesting that immune priming in vaginal challenge may occur faster than in rectal challenge.

The priming models do have some mild technical superiorities over frailty models, namely: (1) in the context of net survival curves, priming models are more general than frailty models, at least in one sense (see the SI); (2) they have strictly concave log-likelihoods (see the SI), slightly simplifying numerical computations; and (3) they nest the reduced model in the interior of their parameter space (see the Materials and Methods section), thereby satisfying the usual hypotheses for theorems about the LRT [50]. Table 3 shows that for most trials, however, if one full model achieves statistical significance at *p*<0.05, several others often follow. Typically, the underlying SIV-macaque data are sparse, and the net survival curves are correspondingly nondescript. If several models show significance, then, no specific biological interpretation can be considered pre-eminent without further investigation.

If some controls remain uninfected after RLCs, e.g., it suggests the confounding hypothesis that the controls may have an unusually low infectability. Fortunately, hyper-challenges with increased viral doses after the RLCs can remove the doubt about low infectability [16, 31, 33], although its cause, immunization or variable frailty, would remain unclear.

To summarize, then, the present article updated Regoes’ database [35] with the SIV-Macaque database in the SI. The updated database displays several trials where the controls appear not to have a constant hazard of infection, and where full models of immune priming (or tolerance) or variable frailty fit control data significantly better at uncorrected *p*<0.05 than the reduced model positing a constant probability of infection per challenge. Given sparse animal data, exact tests have advantages over large-sample tests like the standard logrank test that use Gaussian approximations. Unfortunately, the present results show that any statistical test under consideration should also be shown to be robust against confounding hypotheses about control animals [54], notably both repeated low-dose immunization and variations in frailty. Biologically, the data from SIV-macaque trials are consistent with the RLC protocol changing the infectability of animals after about two weeks in rectal or oral challenge, and possibly even earlier in vaginal challenge. Explanations other than immunization remain very possible, however, particularly because changes in infectability were sometimes consistent with immune priming, and sometimes with immune tolerance. Nonetheless, the results have some bearing on previous biological conclusions that repeated exposure to HIV without treatment may have no effect [35] or may even reduce human infectability [53].

## Materials and Methods

### The SIV-Macaque database

Our meta-analysis [45] permitted an article on an SIV-macaque trial to contribute data to the SIV-Macaque database if it had: (1) at least 5 control animals; (2) at least 3 repeats of the same challenge; (3) a regular challenge schedule; (4) infection assigned to a specific challenge; and (5) publicly available data. Regoes [35] used more stringent criteria than ours, including a restriction to rectal challenges, so his database has 7 articles [12–16, 19, 20] and is a subset of ours. Our SIV-Macaque database includes 2 articles predating his; 2 articles contemporary with his; and 13 articles postdating his.

Table 1 in the Results section displays metadata for our SIV-Macaque database. To favor completeness, our database included two borderline cases: (1) an article exploring the RLC protocol with control animals only [10]; and (2) an article using (almost regular) twice-weekly challenges [17]. The present article followed Regoes [35] in including the Wilson (2009) trial [16], which followed its RLCs with viral hyper-challenges at greatly increased doses to test the infectability of animals uninfected after Challenge 5. As in [35], our SIV-Macaque database includes data corresponding to the RLCs, truncating the challenge data after the RLCs and before any hyper-challenges. The Eudailey (2018) trial [31] also used hyper-challenges after Challenge 8. The Om (2020) trial [33] increased the challenge after Challenge 3, possibly just readjusting the challenge ensure infection of controls, rather than testing their infectability with hyper-challenges.

A reanalysis in the Letvin (2011) trial [19] regrouped some of its data to refine biological conclusions about the *Mamu*-*A*01* allele. Similarly, the Om (2020) trial [33] regrouped their data by sex to refine biological conclusions about vaccine protection by sex. The present article avoided analyzing data twice, so it ignored the regroupings.

Each article yielded challenge data for its trial arms. If the data were figures with net survival curves, the Windows^®^ 10 Snip & Sketch tool captured the curves with a screen shot. Where possible, two people independently digitized each net survival curve with the WebPlotDigitizer (version 4.4) at https://apps.automeris.io/wpd/, later reconciling any discrepancies. No other data format required digitization.

Some articles contained more than one set of control data. Within the articles, however, every arm of every trial mapped to precisely one set of control data. All trial data in an article were grouped by their control arm, and each group was stored as a DataFrame in a separate file of comma-separated values (CSV), following the precepts of tidy data [46]. Each filename corresponded to a composite key combining the article ID in PubMed and the source graphic(s) or table(s) containing its “Control”, anchoring all data unambiguously into the public literature. In the Results section, the columns “PMID” and “Source” in each table implicitly display the composite key for the relevant control.

The next sections detail the various representations used to assemble, retrieve, and analyze the SIV-Macaque database. Flexible representation was indispensable in reducing human effort.

### Data Representation by Individuals

To focus the following discourse, unless stated otherwise, consider only the control arm of an RLC protocol in a specific trial. Following Regoes’ notation [35] where feasible, let *n* = *n*_1_ count the controls, initially uninfected and indexed by *j* ∈ {1,…, *n*}. Each Animal *j* receives serial challenges at regular intervals, e.g., weekly. The challenges are numbered consecutively *t* ∈ {1, 2, 3,…}. Let Challenge *c*_*j*_ be the final challenge to Animal *j*, with *m* = max_*j*_ *c*_*j*_ the largest challenge index among the controls, so *c*_*j*_ ≤ *m*. To account for frequent right-censoring in general survival analysis [44], Regoes [35] carefully permitted the possibility that the final Challenge *c*_*j*_ does not infect Animal *j*, even if *c*_*j*_ < *m*.

On one hand, if Animal *j* becomes infected, Criterion (4) above specifies that the index *c*_*j*_ of the infecting challenge is known. Some trials continue to challenge infected animals, but for present purposes, we ignore such challenges and consider *c*_*j*_ as the final challenge to Animal *j*. On the other hand, the final Challenge *c*_*j*_ may not infect, so some animals may remain uninfected at the end of the trial.

Consider, therefore, the final status (**c**, **i**) = {(*c*_*j*_, *i*_*j*_) : *j* = 1,…, *n*} of every Animal *j*, where *i*_*j*_ indicates whether Challenge *c*_*j*_ infected Animal *j* : *i*_*j*_ = 1 indicates infection, and *i*_*j*_ = 0 otherwise. For a generic animal, drop the subscript *j*, so (*c*, *i*) denotes its final status. The final status (**c**, **i**) is particularly useful if an analysis uses multiple covariates (e.g., alleles) to distinguish individual animals, a rare circumstance indeed in sparse animal trials. We therefore summarize the final status (**c**, **i**) with the statistics relevant to discrete-time hazards in analysis of net survival curves [44].

### Hazards in Discrete Time

Given sampled Animals *j* ∈ {1,…, *n*}, assume a probability model where the corresponding components (*c*_*j*_, *i*_*j*_) of the final status (**c**, **i**) are mutually independent. The hazard *h*_*t*_ gives the probability of infection of a randomly sampled control at Challenge *t*, conditional on previous challenges not having infected it. The Iverson bracket [*A*] provides a standard notation for the indicator of the event *A* : [*A*] = 1 if the event *A* occurs and [*A*] = 0 otherwise [47]. The probability that a generic control has final status (*c*, 0) is (1 − *h*_1_)…(1 − *h*_*c*−1_)(1 − *h*_*c*_) ; final status (*c*,1), (1 − *h*_1_)…(1 − *h*_*c*−1_) *h*_*c*_. In either case, the natural logarithm of the probability that a particular control has final status (*c*, *i*) is

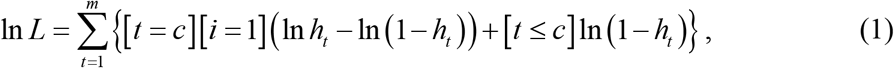

proved as follows. The terms on the right side are 0 for any Challenge *t* after Challenge *c*, so the sum on the right may be broken into terms for *t* = 1,…, *c* −1 and a term for *t* = *c*. Enumeration of two cases [*i* = 0] and [*i* = 1] then shows that Eq (1) gives the natural logarithm of the stated probability.

Let 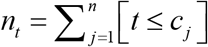 count the controls remaining in the trial at least to Challenge *t*; 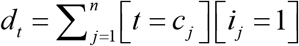, the controls infected by Challenge *t*. The sum over all controls for log-likelihood for the final status (**c**, **i**) in Eq (1) can therefore be rewritten as

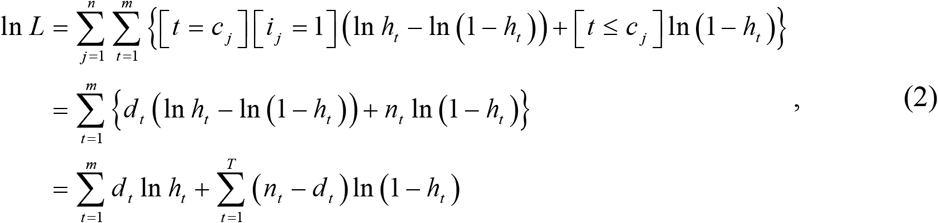

the standard expression from Kaplan-Meier survival analysis [44], put into the present context.

The representation by deaths {(*n*_*t*_, *d*_*t*_) : *t* = 1,…, *m*} quickly yields a net survival curve (e.g., Figure 3 of [18] or Fig 1 below). On one hand, Kaplan-Meier survival analysis derives its generality from handling right-censoring due to incomplete patient follow-up. On the other hand, the RLC protocol in animal trials typically makes follow-up compulsory, so every uninfected control receives its next scheduled challenge. Challenge *t* then infects *d*_*t*_ = *n*_*t*_ − *n*_*t*+1_ of the *n*_*t*_ controls uninfected before Challenge *t* (*t* = 1,…, *m*), where *n*_*m*+1_ counts the uninfected controls at the end of the trial (possibly, *n*_*m*+1_ = 0 ). At the end of a typical trial, each of the *n*_*m*+1_ uninfected controls have therefore survived *m* challenges. Assume below that animal trials have compulsory follow up within each arm, yielding several compact alternatives to the representation by deaths {(*n*_*t*_, *d*_*t*_ ) : *t* = 1,…, *m*}.

**Fig 1.**
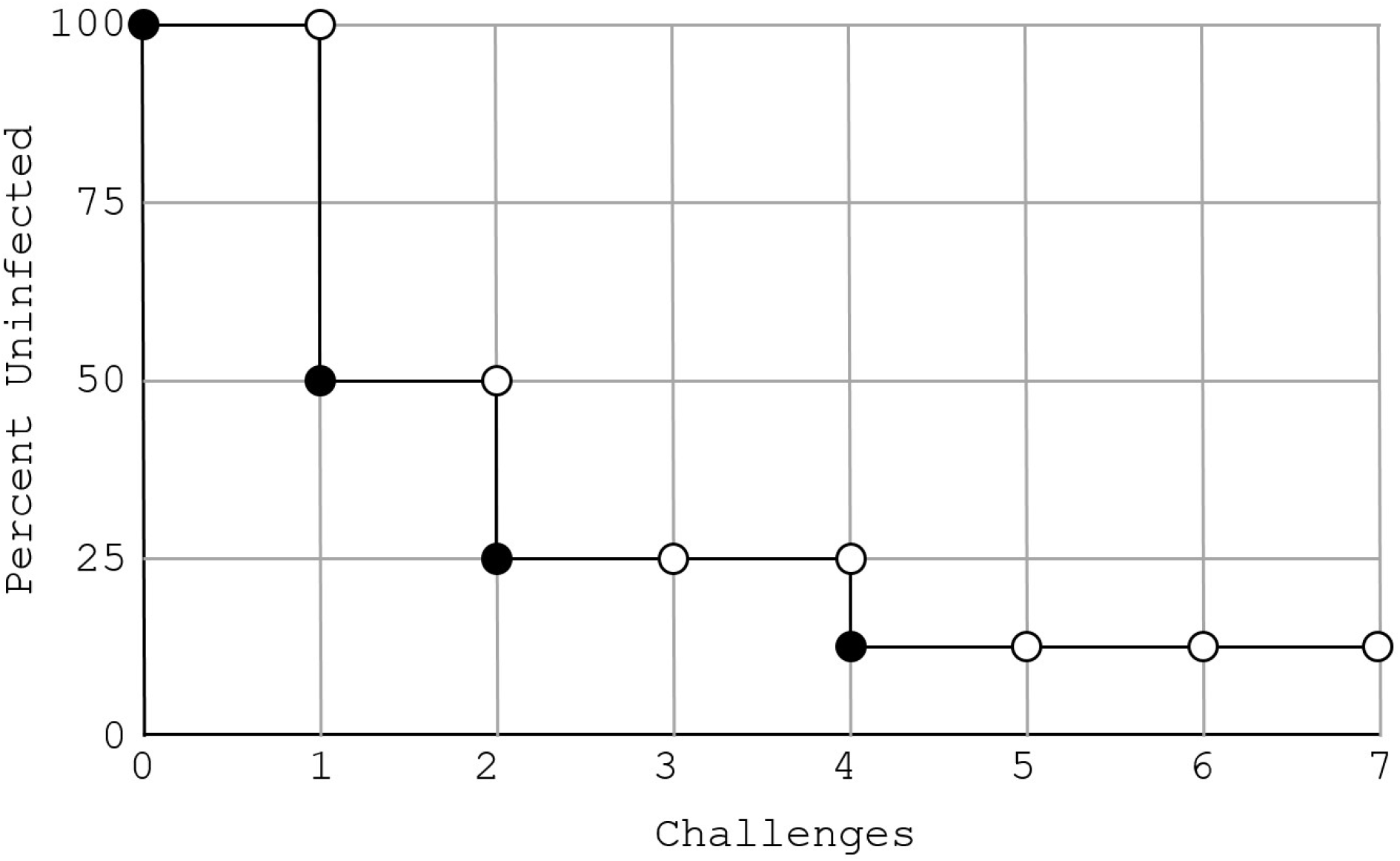
A hypothetical net survival curve for a control arm with 8 animals.

### Alternative Data Representations

(see Fig 1). The representation by survivors (*n*_1_, *n*_2_,…, *n*_*m*+1_ ) satisfies *n*_1_ ≥ *n*_2_ ≥ … ≥ *n*_*m*+1_, and it permits challenge data from each arm in a trial to occupy one column in a DataFrame. The SIV-Macaque database in the SI is a directory of CSV files, each a DataFrame for one trial. Most trials enforce the same maximum *m* of challenges for each arm, so most of the DataFrames are well-formed, with *m* rows in every column and 0s padding an arm after infection of all animals in it. In fact, contrary to an assertion of plurality elsewhere [42], only one trial in the SIV-Macaque database, Ellenberger (2006) [12], overtly permitted *m* to vary with the arm. In the one trial [12], challenge was discontinued before complete infection of controls and before the end of the trial. The corresponding DataFrame therefore has NaNs padding the control arm, instead of 0s.

Fig 1 plots percent uninfected animals against the number of challenges for a control arm with 8 animals. Typical net survival curves use percentages to facilitate comparison of results from trial arms with differing numbers of animals. At each Challenge *t* = 1,…, *m*, the open point (*t*, *n*_*t*_ ) gives the percentage of the originally uninfected animals receiving Challenge *t* ; the closed point (*t*, *n*_*t*_ − *d*_*t*_ ) drops by *d*_*t*_ due to animals infected by Challenge *t* (if not obscured by an open point when *d*_*t*_ = 0 ). By convention, *n*_*m*+1_ counts the uninfected controls at the end of the trial, e.g., *n*_*m*+1_ =1 in Fig 1. Fig 1 also shows that *m* =7, so with *n* =8 animals in the arm, the representation by deaths {(*n*_*t*_, *d*_*t*_ ) : *t* = 1,…, *m*} is { (8,4), (4,2), (2,0), (2,1), (1,0), (1,0), (1,0) }. The representation by survivors (*n*_1_, *n*_2_,…, *n*_*m*+1_ ) uses only the open circles and *n*_*m*+1_ : (8,4,2,2,1,1,1,1). The minimal representation { (*τ*(*i* ) −1, *n*_*τ*(*i*)_ ): *i* = 1,…, *k* } uses only the closed points and (*m*, *n*_*m*+1_ ) : { (0,8), (1,4), (2,2), (4,1), (7,1) }.

A minimal representation reduces the human effort in digitizing the net survival curves from SIV-macaque trials. In the representation by survivors, omit challenges where no infections occurred to yield (*n*_*τ*(1)_, *n*_*τ*(2)_,…, *n*_*τ*(*k* −1)_, *n*_*τ*(*k*)_), where *n*_1_ = *n*_*τ*(1)_ > *n*_*τ*(2)_ > … > *n*_*τ*(*k*−1)_ ≥ *n*_*τ*(*k*)_ = *n*_*m*+1_. Formally, start with *τ*(1) = 1 and define *τ*(*i* + 1) = min {*t* > *τ*(*i*) : *n*_*τ*(*i*)_ > *n*_*t*_ for *t* ≤ *m*} recursively until the set is empty, say, after *τ*(*k* −1) has been defined. Then, define *τ*(*k*) = *m* +1. The minimal representation {(*τ*(*i*) − 1, *n*_*τ*(*i*)_ ): *i* = 1,…, *k*} compacts the representation by survivors without loss and greatly speeds digitization of net survival curves. Most experiments show the net survival curve for each arm as percent survival, as in Fig 1. Accordingly, we digitized only the points corresponding to the minimal representation, multiplied the percentages by *n* = *n*_1_, and then rounded to integers to yield the minimal representation. If two people digitize and then disagree, the minimal representation facilitates a rapid reconciliation of their results.

Finally, a few trials (e.g., [13, 16]) use a representation (*t*_*j*_ : *j* = 1,…, *n* ) by individual infecting challenges [35], where Challenge *t*_*j*_ infects Animal *j*. Retain our convention that *t*_*j*_ = *m* + 1 if Animal *j* remains uninfected throughout the trial. Then, the data may be converted to *n*_*t*_ = # {*t*_*j*_ ≥ *t* : *j* = 1, 2,…, *n*} for *t* = 1,…, *m*, where “ # ” counts the number of elements in the set.

### Constant Hazard reduced model

The following nests a reduced model *H*_0_, the Constant Hazard model, within several different full models, to detect deviation from *H*_0_ with Likelihood Ratio tests (LRTs). The Constant Hazard model (Regoes’ Geometric infection model [35]) posits that all challenges yield mutually independent results and have a constant probability *p* of infecting a control. Every animal is equally infectable, so under the reduced model *H*_0_, the hazard per challenge is the constant

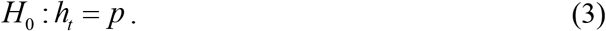

The corresponding log-likelihood

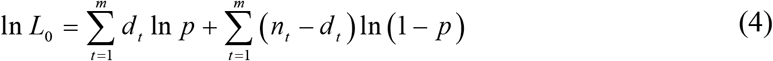

yields the maximum likelihood estimator (MLE)

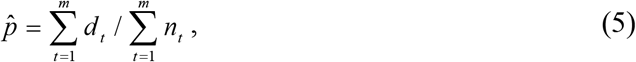

a convenient initialization for maximizing likelihoods of the full models. Even if the reduced model is rejected, Eq (5) is still the fraction of all challenges that infected and provides a practical summary of the average potency of a challenge.

For brevity, to truncate a real number *x* to a probability *U*(*x*) between 0 and 1, define the unit function *U*(*x*) = *x* for 0 ≤ *x* ≤ 1, with *U*(*x*) = 0 for *x* < 0 and *U*(*x*) = 1 for *x* > 1. Below, we characterize several full models *H*_•_, where “ • ” is an arbitrary subscript, simply by specifying their hazard *h*_*t*_. If required, Eq (2) then implicitly gives the corresponding likelihoods *L*_•_ and log-likelihoods *ℓ*_•_ = ln *L*_•_. Our full models include all of Regoes’ full models [35].

The first elaboration of the Constant Hazard model that Regoes [35] addressed was immune priming: even if the animals had the same infectability, their infectability might evolve during the trial because the RLCs provided immunization against the virus, either in the direction of immune priming or immune tolerance. In mathematical models, negative immune priming corresponds to immune tolerance.

### Arithmetic priming full model (AP)

Regoes [35] examined a full model equivalent to

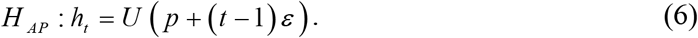

*H*_*AP*_ reduces to *H*_0_ if *ε* = 0. Priming corresponds to *ε* < 0 ; tolerance, to *ε* > 0.

### Geometric priming full model (GP)

Arithmetic priming has a hard boundary at 0 from the unit function. A geometric sequence avoids it:

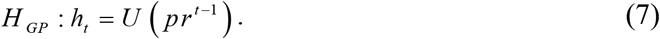

*H*_*GP*_ reduces to *H*_0_ if *r* = 1. Priming corresponds to *r* < 1 ; tolerance, to *r* > 1.

### Step priming full model (SP1, SP2, SP3)

Following Regoes [35] (see also [48]), consider full models where the hazard is piecewise constant, changing from *p* to *p*_2_ after Challenge *l*_*step*_ =1,2, or 3.

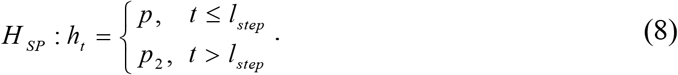

*H*_*SP*_ reduces to *H*_0_ if *p*_2_ = *p*. Priming corresponds to *p*_2_ < *p* ; tolerance, to *p*_2_ > *p*.

The log-likelihoods for priming models are all strictly concave in their two variables ( *p*,*ε*), ( *p*, *r* ), or ( *p*, *p*_2_ ) (see the SI), so hill-climbing methods always determine the MLE as the unique global maximum. Regoes [35] also addressed frailty models without priming, where animals might have different intrinsic infectabilities that remained constant during challenge [49]. The log-likelihoods for the two frailty models below are not concave in general (see the SI), mildly complicating numerical computation. In addition, the two frailty models nest the reduced model on the boundary of their parameter space, violating the hypotheses of standard statistical theorems [50]. The present article has the purview of examining the robustness of Regoes’ scientific conclusions against expanding his database, making technically arduous statistical improvements peripheral.

### Beta Frailty full model (BF)

The Beta distribution with parameters (*a*, *b*) has probability density function

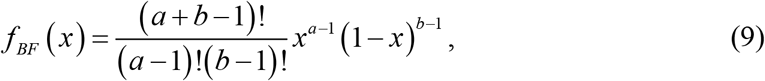

where factorials replace the cumbrous Gamma function notation, even if *a* and *b* are not integers. The prefactor is the reciprocal of the Beta function

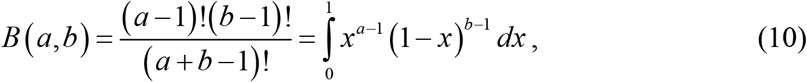

yielding the required normalization 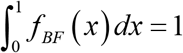.

Consider a frailty model where each challenge to Animal *j* has a probability *x* of infection, the infectability *x* being intrinsic to Animal *j* with the probability density function in Eq (9). The corresponding hazard, the probability of infection on Challenge *t* conditional on previous non-infection, is

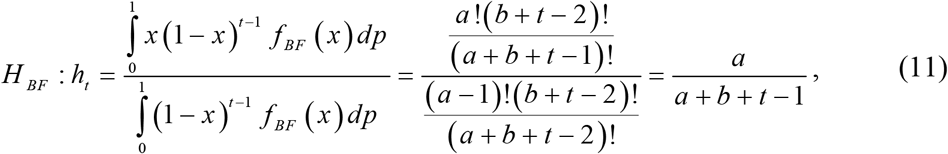

equivalent to animal frailty with a beta distribution (see the SI). Hence, the log-likelihood

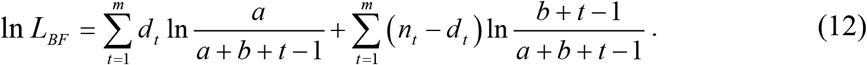

To nest *H*_0_ within *H*_*BF*_, Regoes [35] noted that the beta distribution in Eq (9) can be parametrized by its mean *p* = *a* / (*a* + *b* ) and variance *v* = *ab* / (*a* + *b* )^2^ / (*a* + *b* + 1). For *v* > 0, let *s* = *p* (1 − *p* ) / *v* − 1 so *a* = *sp*, *b* = *s* (1 − *p* ). *H*_*BF*_ reduces to *H*_0_ if *v* ↓ 0 with *p* fixed, i.e., *a*, *b* → ∞ with *p* = *a* / (*a* + *b* ) fixed.

### Delta Frailty full model (DF) [34, 49]

In the context of a treatment trial, Hudgens et al [34] posit a frailty model for controls with parameters (*θ*, *p*). In their model, each control is independently susceptible with probability 1 − *θ*. Only susceptible controls can be infected, and each challenge infects susceptible controls with probability *p*. The analog to Eq (9) is

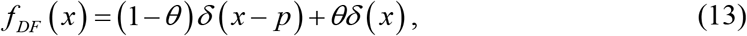

where *δ*(*x*) symbolizes the Dirac delta, which satisfies 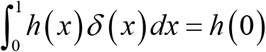 for functions *h*.

Thus,

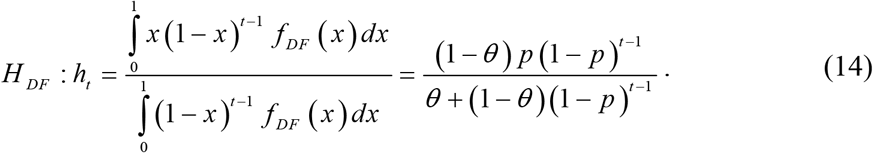

*H*_*DF*_ reduces to *H*_0_ if *θ* = 0.

### Model Parameter Estimation

Animal data are sparse, so if the reduced model holds, we base a confidence interval for *p* on likelihood ratios [35, 51], rather than moments.

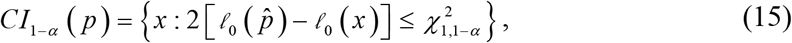

where *α* is the desired significance level and 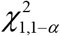 is the (1 − *α*)-percentile of the chi-squared distribution with one degree of freedom, i.e., for a random variate *X* with the distribution, 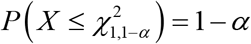. We take *α* =0.05 to yield 95% confidence intervals. To facilitate simulation of actual trials, the Excel file in the SI contains the MLE for each full model, along with the corresponding Fisher information matrix, if calculable numerically.

## Acknowledgments

I acknowledge the invaluable aid of Ms. Chelsea Zebaze in digitizing data from animal trials and several useful conversations with Dr DoHwan Park and Dr Junyong Park.

## Supporting information

**Supplementary Information.zip. This is a ZIP file of the following items. README.txt. This is an index for the Supporting Information.**

**Supplementary_Methods.docx. This contains additional methods and a legend for Supplementary_Table.xlsx.**

**Supplementary_Table.xlsx. This table includes Tables 1–3 and other information in floating-point precision.**

**SIV-Macaque_Net_Survival_Curves/. This is a directory containing the CSV files, DataFrames containing the data from the net survival curves in the SIV-Macaque database.**

